# Re-analysis of Transcriptomic and Proteomic Data Using Multi-Omics Approaches Identifies Biomarkers of Diabetes-Associated Complications in an INS Mutant Pig Model

**DOI:** 10.64898/2026.03.04.707880

**Authors:** Kota Krishna Priya, Bilal Ahmed Abbasi, Priya Kajla, Sadhana Tripathi, Aubrey Bailey, Binuja Varma

## Abstract

Mutant Insulin Induced Diabetes of Youth (MIDY) is an established porcine model caused by the INSC94Y mutation, which results in misfolded insulin, leading to severe β-cell loss and hyperglycemia. Understanding disease pathophysiology is critical for identifying biomarkers and therapeutic targets, and animal models play a key role in this process. In this study, we re-analyzed published transcriptomic and proteomic data from the MIDY model using advanced multi-omics approaches and our in-house SurfacOmics tool. This integrative analysis identified ADAMTS17 as a novel biomarker, suggesting a potential association in diabetes-associated immune dysfunction and delayed wound healing through ECM-immune interplay.

## Main

Despite extensive research into diabetes-associated molecular pathways, identifying reliable biomarkers to distinguish disease states remains a significant challenge (1). ADAMTS family members have been implicated in extracellular matrix (ECM) remodeling and immune regulation, yet their role in diabetes remains underexplored.

In this work, we revisited a report by Backman et al (2) that builds a transgenic biobank resource model of chronic insulin-deficient pigs (*Sus Scrofa)* (3). This MIDY model was developed by inducing mutagenesis at *INS*^C94Y^ to disrupt disulphide bonds between alpha and beta chains, resulting in misfolded proteins (4). The accumulation of misfolded proinsulin in the ER leads to apoptosis of insulin-producing β-cells (5), Thus, MIDY pigs are severely diabetic with insufficient β-cell mass and require exogenous insulin treatment which mimics poorly controlled diabetic patients. It is important to note that the MIDY model does not fully reflect all disease mechanisms of T1D, as it lacks the autoimmune component inherent to T1D. Backman et al.’s study examines the transcriptome and proteome of a small cohort of 2-year-old female transgenic model animals, MIDY (n=4) and paired wild-type (WT) littermates (n=5).

In our independent re-analysis, we co-examined transcriptomic and proteomic data in an integrated, multi-omics’ context to identify biomarkers associated with diabetes-induced molecular alterations. We downloaded transcriptomic data from the GEO repository (GSE122029) as raw FASTQ files and fetched proteomic LC-MS/MS data from ProteomeXchange consortium (PXD011536). The raw data contained 34328 genes and 2535 proteins. After several rounds of preprocessing and filtering, datasets containing 19,664 genes and 489 proteins passed quality filters and were retained for downstream analyses. (**Supplementary 1.1**).

A preliminary analysis of the filtered proteomic dataset showed a clear separation between WT and MIDY groups demonstrated by PCA (**Figure 1A**). Associated component loadings indicated that the greatest proportion of variance in the combined dataset was attributable to differences in HMGCS2 and RDH16, metabolic enzymes which corroborate the original findings from Backman et al., 2019. Notably, COL6A3, a key ECM component, showed reduced levels in the MIDY group, while ADAMTS17 levels were elevated when compared with reference WT (**Figure 1B-D**). A Pearson correlation coefficient of –0.48 between ADAMTS17 and COL6A3 supports their inverse relationship and suggests that dysregulation of ADAMTS17 may influence collagen composition in diabetic states (**Figure 1B**).

**Figure 1.**
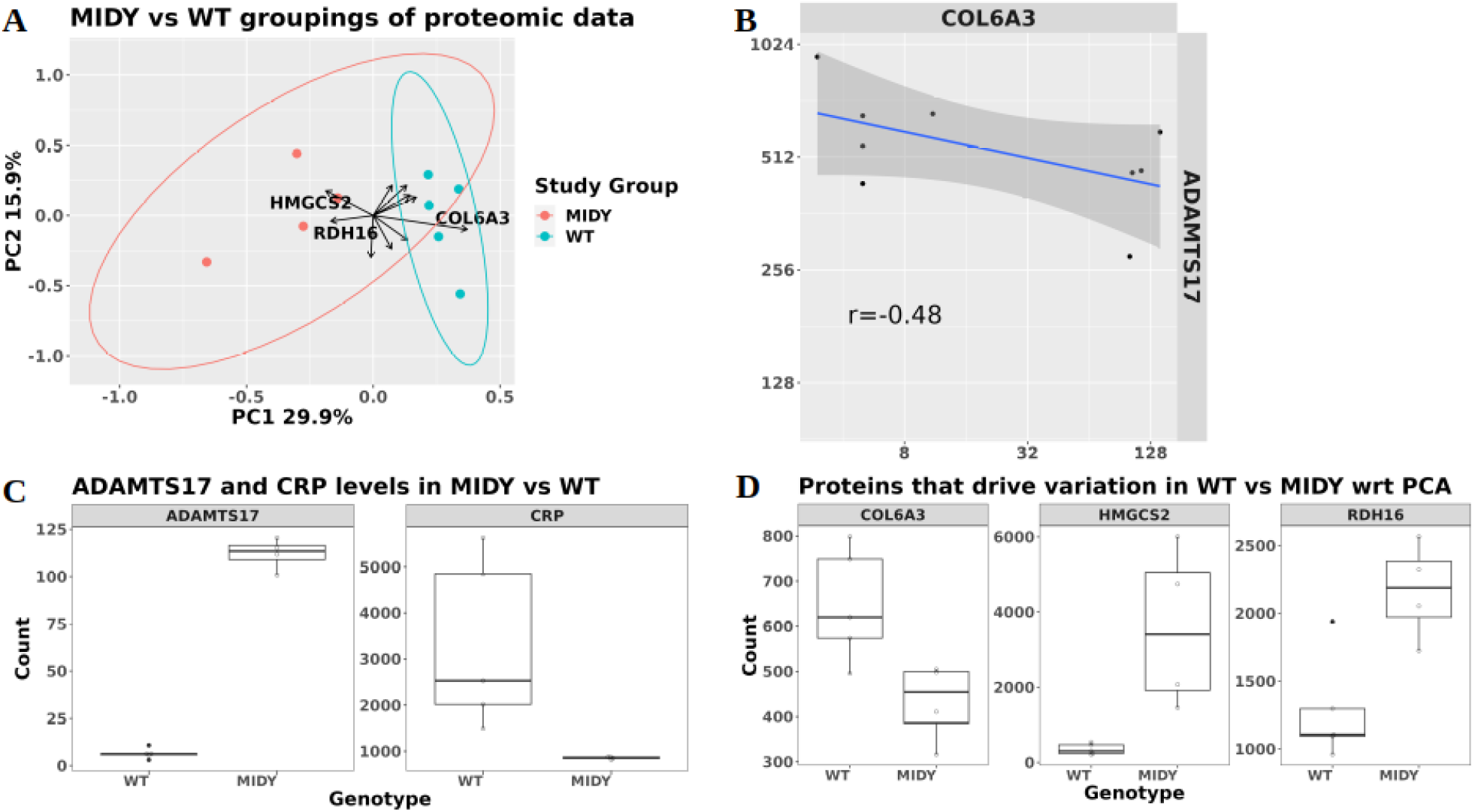
**(A)** PCA plot demonstrating the groupings of WT and MIDY groups at proteome level, COL6A3 highlighted as driver gene for WT group **(B)** Negative correlation between ADAMTS17 and COL6A3 suggesting the role of ADAMTS17 in ECM degradation **(C)** *ADAMTS17* and *CRP* levels in the MIDY vs WT groups. **(D)** Protein levels of PCA driver genes distinguishing WT and MIDY groups

Differential expression analysis of the transcriptomic dataset revealed a further 17 genes with log-fold change > 1 (**Supplementary Table 1**). No differentially abundant proteins were identified at an FDR < 0.01. These findings were also concordant with those reported in Backman et al. 2019. However, we consider that the relevance of changes in COL6A3, ADAMTS17, and CRP **(Figure 1C-D)** may have been under-considered as evidence of some degree of immune incompetence in the MIDY model.

Little is known about the functional relevance of ADAMTS17, apart from its sequence similarity (>30% peptide identity) to ADAMTS4 and ADAMTS5 suggesting a potential role in ECM regulation and immune modulation (**Supplementary Table 1, Supplementary 1.2**). From this familial relationship we can infer its role in reshaping the ECM, and the immune response. Akyol S et al. also reported a role in cartilage proteoglycan degradation in diabetes mellitus patients (6).

Integrated multivariate analyses based on the Partial Least Squares (PLS) framework were performed, including sparse PLS (sPLS) for unsupervised feature and supervised multiblock PLS approaches (DIABLO, mbPLS), using “mixOmics” package (version 6.23.4) (7) on R version 4.3.2(**Supplementary 1.3**). These approaches facilitated the identification of molecular mechanisms underlying chronic insulin deficiency. Both transcriptomic and proteomic integration analyses consistently identified ADAMTS17 (transcriptomic data) and fibrinogen gamma (FGG; proteomic data) as top discriminative features distinguishing MIDY from WT groups (**Supplementary Table 2**).

Functional network analysis using STRINGdb (5% FDR, confidence 0.15) identified two prominent co-expression modules derived from sPLS-selected genes: one representing metabolic and another representing immune signatures **(Figure 2A)**. The immune module included CXCR2, CD40, and CD59, which clustered distinctly from the metabolic genes and exhibited lower expression in MIDY compared to WT reference **(Figure 2B)**. Also, the CRP levels were reported to be significantly reduced in the MIDY group relative to the WT reference, consistent with the observed differences in immune marker expression (**Figure 1C**). **Figure 2B** further confirms this by showing relatively lower levels of these immune markers in MIDY group. This pattern reflects an attenuated immune response in the MIDY group, consistent with chronic insulin deficiency associated with immune suppression.

**Figure 2.**
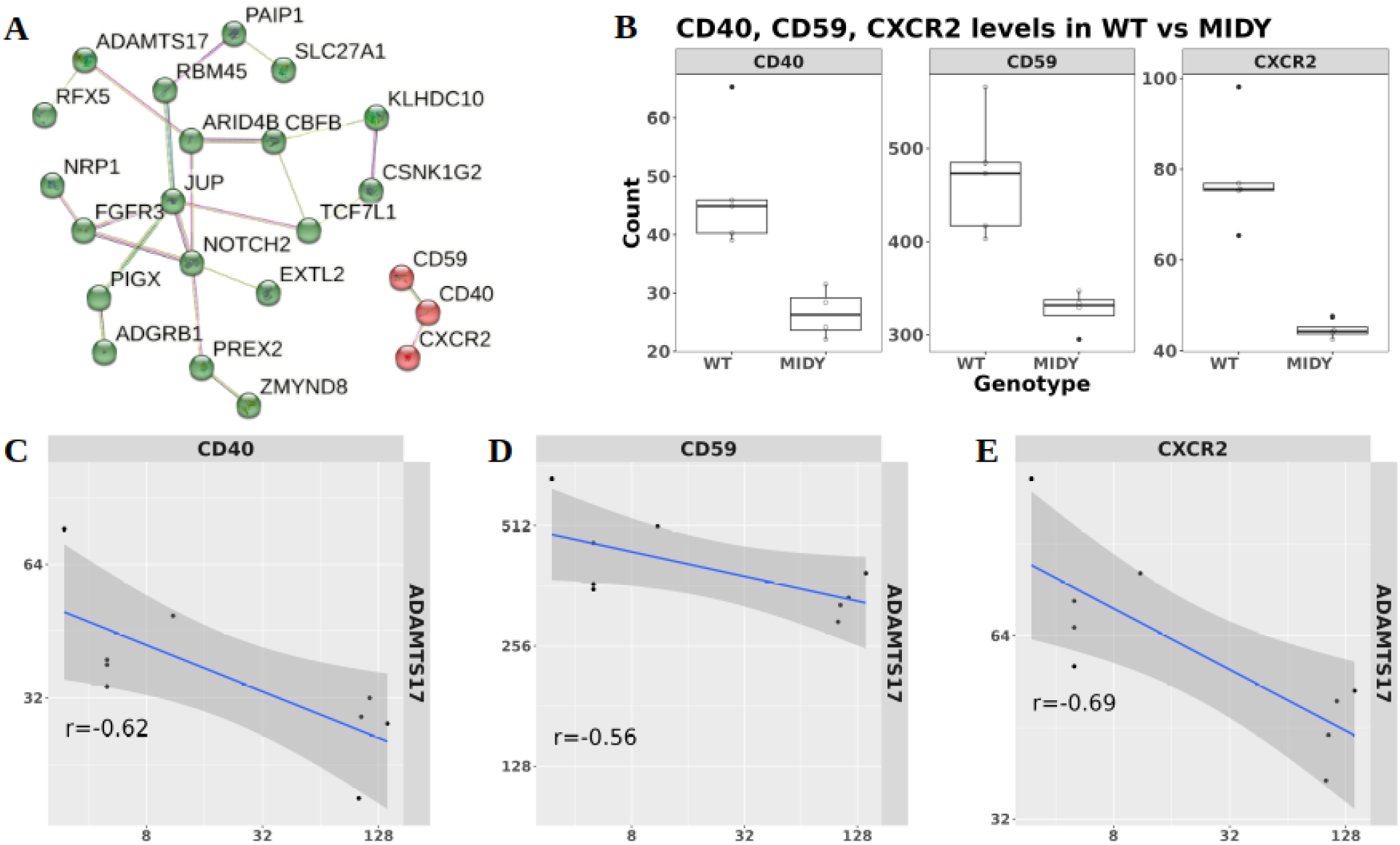
**(A)** STRING network revealing the immune signatures (highlighted in red) and metabolism-centric genes (highlighted in green) selected via sPLS as two separate modules, and **(B)** Levels of CD40, CD59 and CXCR2 in MIDY and WT groups and **(C, D, E)** Negative correlations of CD40, CD59 and CXCR2 immune markers with ADAMTS17 suggesting an association between elevated ADAMTS17 levels and reduced immunerelated marker expression.

Correlation analysis further supported the association between ADAMTS17 and immune markers. A strong negative correlation was observed between CD40, CD59, CXCR2 and ADAMTS17 (**Figure 2C, D, E**), indicating that elevated ADAMTS17 levels may accompany reduced immune signalling. These observations suggest that ADAMTS17 might contribute indirectly to immune modulation, possibly through ECM-cytokine interactions rather than direct enzymatic degradation.

To further strengthen these findings, ADAMTS17 was examined for associated genes using DESeq2 (8) linear correlation analysis. Significant correlations were found with MBL2 (**Supplementary Table 1**), which play a key role in tissue-repair and fibrinogen-thrombospondin complex formation along with FGG. Elevated levels of ADAMTS17, together with MBL2 and FGG, reinforce a possible interplay between ECM remodeling and immune response under diabetic conditions (9).

To evaluate the translational potential of these gene candidates, we applied our in-house developed biomarker discovery R Shiny based application, *SurfacOmics* (10), designed to integrate multi-omics signals with subcellular localization and marker assayability. *SurfacOmics* identified ADAMTS17 as the top transcriptomic biomarker and FGG as the top proteomic biomarker based on marker potential scores (**Figure 3 and Figure 4B, Supplementary 1.4**). Both biomarkers exhibited clear expression differences between MIDY and WT groups (**Figure 3 and Figure 4A**). Notably, ADAMTS17 and FGG demonstrated strong marker potential, by the tool recommending them as assayable targets based on their cellular localization. Despite not being located on the cell surface, FGG has a commercially available polyclonal antibody to detect its levels in the cytoplasm by immunofluorescent assays **(Figure 3-4B)**.

**Figure 3.**
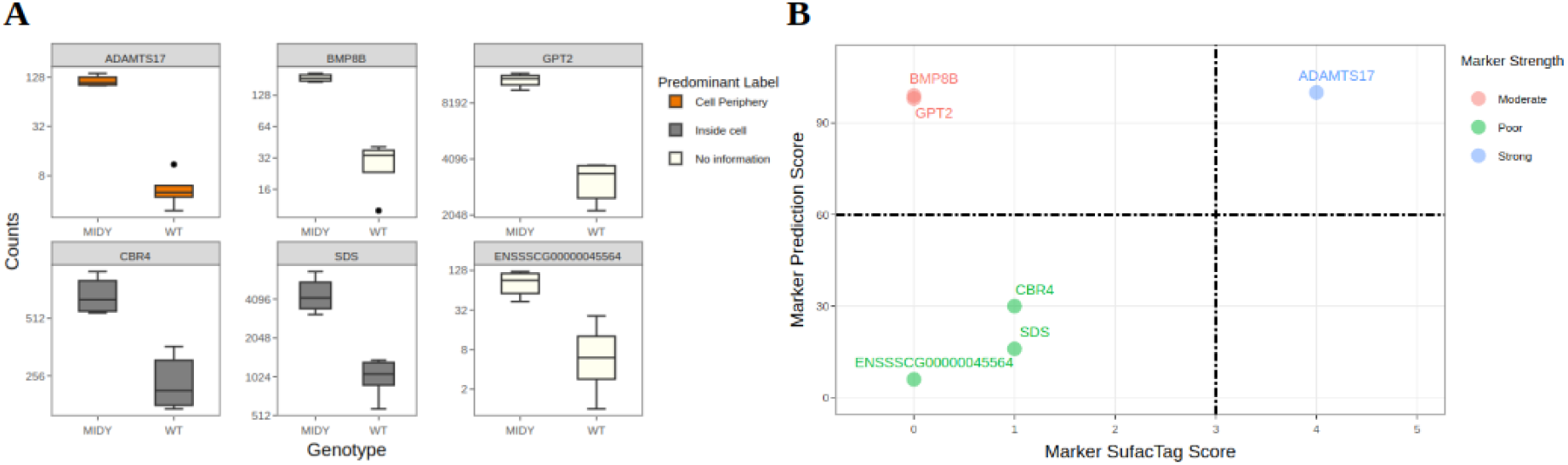
**(A)** Expression difference of *SurfacOmics*-identified transcriptomic biomarkers between the MIDY and WT groups. **(B)** *SurfacOmics* marker potential plot, showing ADAMTS17 (highlighted) as a potential biomarker.

**Figure 4.**
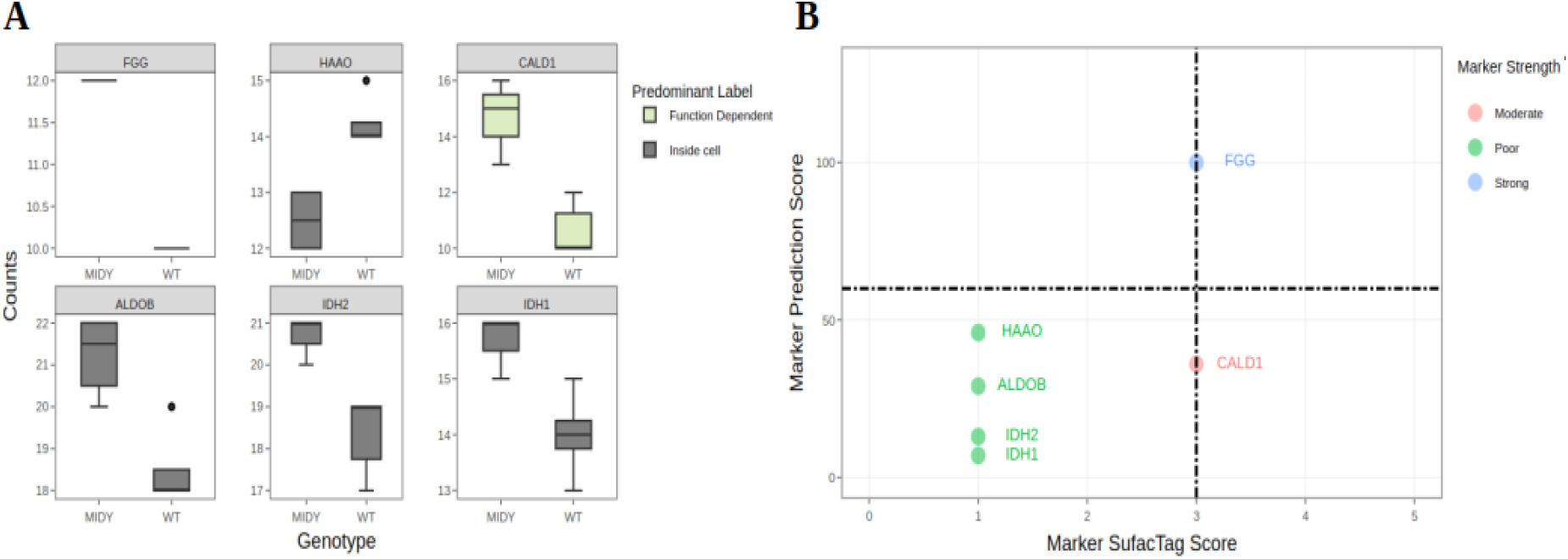
**(A)** Expression difference of *SurfacOmics*-identified proteomic biomarkers between the MIDY and WT groups. **(B)** *SurfacOmics* marker potential plot, showing FGG (highlighted in blue) as a potential biomarker.

In summary, this integrative multiomics reanalysis identifies ADAMTS17 as a consistent molecular signal associated with extracellular matrix remodeling and altered immunerelated marker profiles in the MIDY pig model. The convergence of ADAMTS17 with FGG and MBL2 across multiple analytical frameworks suggests coordinated molecular changes linked to chronic insulin deficiency. While further experimental validation is required, these findings demonstrate the value of integrative computational approaches, including SurfacOmics, for prioritizing candidate biomarkers relevant to diabetes associated molecular alterations.

