## Supplementary Information for "Re-analysis of Transcriptomic and Proteomic Data Using Multi-Omics Approaches Identifies Biomarkers of Diabetes-Associated Complications in an INS Mutant Pig Model"

**SUPPLEMENTARY FILE**

### 1.1 Transcriptomic and Proteomic data processing

The raw FASTQ files were preprocessed by implementing RNA-seq analysis Snakemake workflow (RASflow), and the raw read quality was evaluated using MultiQC. After the quality check, reads were trimmed for adapters using Trimmomatic and then aligned to the *Sus Scrofa 11.1* reference genome using HISAT2. Reads were subsequently counted using featureCounts. The count data was then annotated for gene attributes via BiomartDB and improvised with bioDBnet. This raw count data was normalized, and a custom filtering procedure is applied to remove low quality samples and lowly expressed genes. First, the samples with insufficient sequencing depth were excluded by retaining only those with a total read count exceeding 8 million reads. Size factors were estimated to normalize differences in library size across the samples. Gene-level filtering was subsequently performed on normalized counts, retaining genes with atleast five counts in a minimum of three samples. This filtering strategy reduced technical noise from low-coverage samples and low-abundance transcripts, resulting in 19664 reliably expressed genes.

The fetched MS/MS data of 2535 proteins was initially annotated for standardization using Uniprot database. The peptide count data was collated by merging the peptide isomers and precursors and averaging the counts to the protein/gene level. The average count was calculated for each protein. Proteins with poor counts (0, 1) across the samples were discarded. This resulted in a 2475 protein data set. These proteins were processed using a filtering strategy to remove low-abundance and sparsely detected proteins. Size factors were first estimated to account for global differences in sample-wise signal intensity. Proteins were then filtered, retaining only those with raw counts of atleast five in a minimum of six samples. Following this, an additional abundance-based filter was applied, whereby proteins with a mean raw count below 10 across all samples were excluded. This two-step filtering procedure reduced noise and resulted in 489 proteins with sufficient and reliable quantitative signal. These 489 proteins were annotated to the gene level and have been used for all downstream analyses.

### 1.2 Multiple Sequence Alignment (MSA) of ADAMTS17 and its family members having a known role in Diabetes Mellitus

MSA was performed to evaluate the similarity among ADAMTS17, ADAMTS4 and ADAMTS5 protein sequences via Clustal Omega (https://www.ebi.ac.uk/Tools/msa/clustalo/).

### 1.3 Multi-Omics analyses of transcriptomic and proteomic data

Multivariate analyses were performed to identify the distinguishing features between MIDY and WT. To establish the correlations between RNA and protein genes, we implemented sPLS (sparse partial least squares), an unsupervised technique from the "mixOmics" package (version 6.23.4). To identify the crucial genes, we performed component tuning (up to 10-fold cross-validation on top 5, five repeats) followed by variable tuning (10-fold cross-validation, five repeats, on 19664 and 489 transcriptomic and proteomic features respectively). Variables selected were used to construct the plots.

We further checked the correlations using two supervised multi-block PLS-DA algorithms called the DIABLO framework from the "mixOmics" package and Python-based mB-PLS (multiblock PLS) on transcriptomic and proteomic data. For DIABLO analysis, component tuning was performed via a 4-fold cross-validation for 10 repeats on three components. Then, variable tuning was performed on 19664 transcriptomic and 489 proteomic features for all samples using 4-fold cross-validation for 10 repeats. MBPLS was employed to gain deeper insights into the relationships and patterns across different data sources and their associations with the response variables of interest**.** LOO-CV MSE (Leave one-out cross validation Mean Squared Error) plot is used to pick the initial no. of components to test. After testing the variance explained by each component, we picked only the first component as it explained 95.84% variance in the dataset for both RNA and Protein blocks. Then we sorted the features and picked the top 15 "positive" and bottom 15 "negative" discriminative loadings (features) from each block.

#### 1.4 Identification of biomarkers via in-house *SurfacOmics* application

SurfacOmics uses elastic-net penalization combined with a scoring system along with a manually curated "SurfacTag" knowledgebase designed to identify and prioritize surface markers, provided the metadata, count data and input genes/proteins of interest. *SurfacOmics* assigns two scores for each identified marker. Primarily a Marker Prediction Score (MPS), which explains how well the identified marker distinguishes MIDY and WT groups. Secondly, a Marker SurfacTag Score (MSS), which specifies the cellular localization of the identified marker. If the identified marker is readily accessible on the surface and easily assayable, the score will be higher, and vice versa.

The existing version is applicable for annotated transcriptomics and proteomics for six different organisms (*Homo sapiens, Mus musculus, Sus scrofa, Drosophila melanogaster, Macaca mulatta, and Danio rerio*).

### Supplementary Tables

Supplementary Table 1. DEGs identified in WT vs MIDY samples, Multiple Sequence Alignment similarities among ADAMTS17, 4 and 5. ADAMTS17 associated genes from DESeq2 linear correlation analysis.

Supplementary Table 2. Transcriptomic and proteomic features identified by various multivariate techniques.
